# Allocentric Flocking

**DOI:** 10.1101/2025.01.27.634610

**Authors:** Mohammad Salahshour, Iain D. Couzin

## Abstract

Understanding how group-level dynamics arise from individual interactions remains a core challenge in collective behavior research. Traditional models assume that animals follow simple behavioral rules, like explicitly aligning with neighbors, yet experimental support for such interactions is often lacking. Here we consider a model grounded in the neurobiological principles underlying animals’ navigational circuits, particularly the fact that animals encode their headings, and also bearings to objects (e.g., other individuals) in their environment, via a world-centered—allocentric—neural coding. We compare this to an egocentric representation, where bearings are encoded with respect to the arbitrary heading of the animal. An allocentric framework, as opposed to an egocentric one, is shown to enable effective tracking of dynamically moving targets. Moreover, we demonstrate that when individuals themselves act as sensory inputs to each other, that sophisticated, coherent collective motion can emerge directly from navigational circuits (and thus, may readily evolve in nature), without requiring explicit alignment, or additional rules of interaction.

## Introduction

How collective behavior arises from interactions among individuals is central to multiple scientific disciplines [1–4]. A particularly notable example is collective motion; beyond its aesthetic appeal, collective motion has been a testing ground for theories of collective behavior [5]. This is because the emergent macroscopic patterns arise from feedback between the individuals and the collective [6, 7]. Traditionally, models of collective movement were rooted in agents following simple behavioral rules. Such studies have shown that emergent patterns can arise among such cognitively minimalist agents, termed ‘self-propelled particles’. While the earliest such models included explicit alignment—such as the influential Vicsek model [8]—other models have shown that collective motion can arise from mechanisms like escape and pursuit [9], inelastic collisions [10, 11], attractive and repulsive radial forces [12–18], active elastic forces [19, 20], and nematic collisions [21], all of which can induce local alignment.

While suitable for inanimate objects or simple organisms, these modeling frameworks overlook the cognitive processes that shape individuals’ perception of their physical and social environment [22–25]. This realization has led to more recent models that incorporate mechanisms like visual sensing of neighbors [22, 26–28], and the explicit consideration of the sensory-motor interface, such as by incorporating biologically-plausible mechanisms by which individuals may modify both their movements, and their internal model of the world [29–33].

Despite these advancements, however, the relationship between known neural architectures for translating sensory information to movement remains poorly understood in the context of collective behavior. To address this gap, here we present a model that considers the way in which the brain of both invertebrates [34, 35] and vertebrates [36] encodes and processes spatial information when making movement decisions. A central feature of this process is the neural representation of inanimate targets [37–39] and/or conspecifics [34, 35, 40], as well as the required consensus dynamics on ring-attractor neural networks that establish the resulting “decision” in the form of a “neural bump” that influences future movement.

Ring attractor networks, the neuroanatomy of which has been described in insects [34, 35] and in fish [36], and whose phenomenology has been found in mammals [37–39, 41], play a central role in spatial navigation. For example, they have been employed to account for diverse phenomena ranging from working memory [42] and head direction [43–49] to view cells [50], and grid cells [51]. Motivated by the vectorial representation of space in the brain [35, 37–39], it has recently been shown that ring attractor networks also account for how fruit flies, locusts and zebrafish (and thus both invertebrates and a vertebrate) make decisions when choosing between multiple spatial “targets” [40,52,53]. Targets, such as conspecifics, are represented as neural activity bumps on the ring, with ring position relating to corresponding perceived directions to the targets, and computation on the ring occurring via local excitation and long-range/global inhibition. The timescale of the neural dynamics, being typically faster than the movement decisions, results in spatial decisions being considered as an “embodied” process, whereby the movement of the focal animal (or conspecific(s)) changes the geometrical neural representation, which changes the resulting neural bump, that encodes the direction, and speed, of subsequent motion, which changes the geometry of the representation, and so on.

In the context of collective behavior, however, while the ring-attractor framework was found to explain accurately the response of an individual responding to one, two, or three conspecifics [40, 53], it could not account for the consistent coherent motion seen in mobile animal groups (unless explicit local alignment is reintroduced [28], thus divorcing the model from neural principles and experimental data [54, 55]).

Further to this, irrespective of their differences, all models of individual and collective motion (including the widely-used rule-based zonal models [12–14, 56],) make a universal assumption; that vectorial information regarding conspecifics (the estimated directions/bearings towards others) are considered exclusively from an egocentric perspective. That is, it has always been assumed that, with respect to conspecific bearings, the frame of reference for a focal individual is with respect to its own, present, heading (for example, a neighbor positioned directly to the right would be at +90 degrees, whereas one at the left would be considered -90 degrees, with respect to the focal individual’s heading, irrespective of its absolute heading, i.e., whether it is traveling N, S, E or W, see Fig. 1).

**Figure 1:**
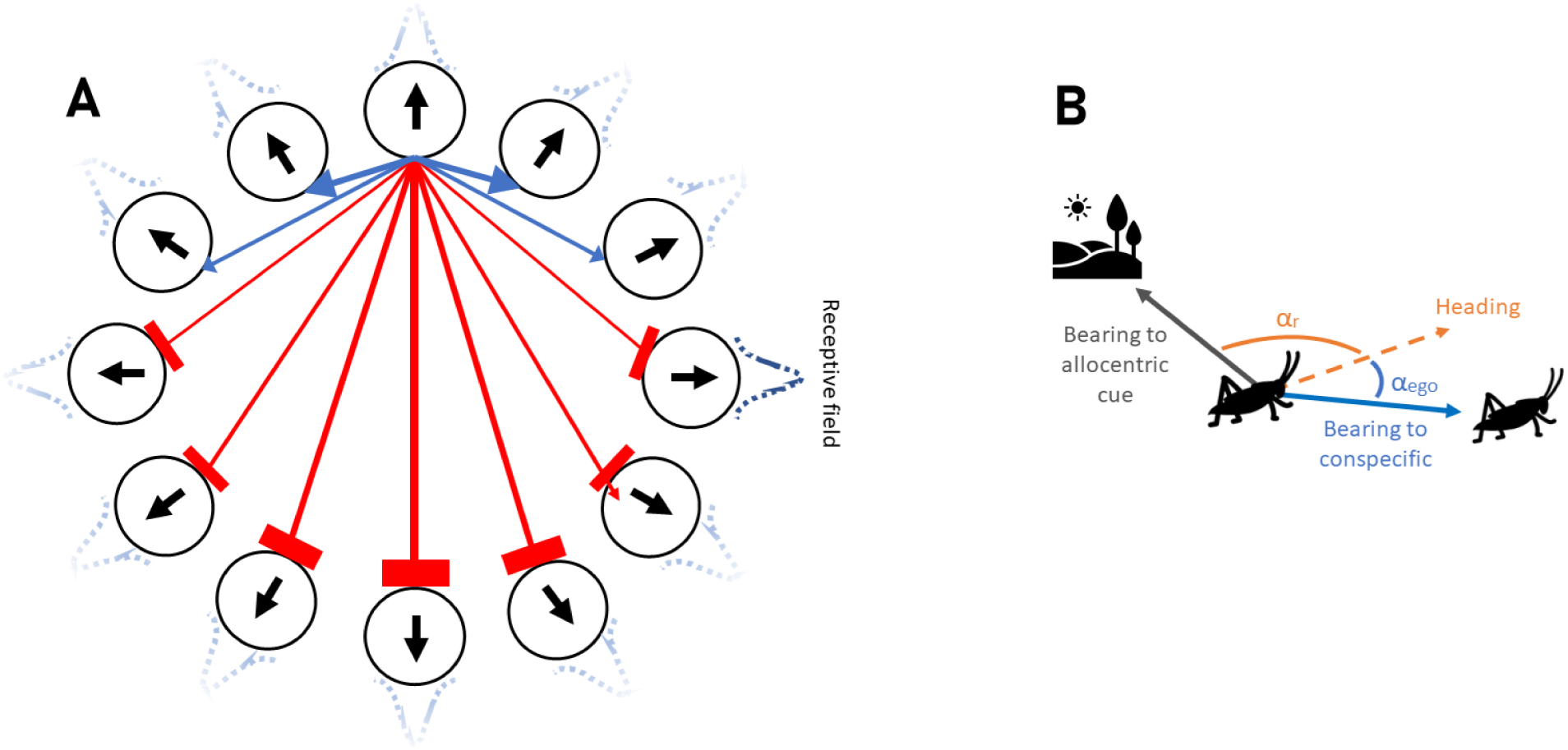
Ring attractor networks with an allocentric and an egocentric perception of space. **A**: Individuals are equipped with a ring attractor network in which neurons are arranged on a ring. Each neuron group, modeled as a spin variable, receives input from the external world through a Gaussian receptive field and induces a movement along the same direction once it is active. Besides, neuron groups interact with other neuron groups via excitatory or inhibitory synapses depending on their distance along the ring. **B**: With an egocentric representation of space, the animal encodes directions with respect to a self-body coordinate (head direction), α*_ego_*. Whereas, with an allocentric representation, directions towards targets are encoded via an allocentric frame of reference, α*_allo_* = α*_r_* + α*_ego_*, where α*_r_* is the direction of the allocentric frame of reference. Thus, directions are independent of the agent’s body coordinate.

By contrast, however, neurobiological data demonstrate that the bearing towards external goals, even in simple animals such as the fruit fly (*Drosophila* species), is encoded in an allocentric (i.e, world-centered, such as north, south, east, west) frame of reference, and that such a representation is ubiquitous among animals [57–62]. In the best-studied example of this, *Drosophila*, it is clear that the brain has evolved to utilize a wide range of cues, such as polarized light [63, 64] and/or prominent relatively stationary objects in the environment (including the sun [65], or landmarks [66], and in some species possibly magnetic fields [66, 67]), to allow the fly to rotate the cellular activity in it its neural compass (also a ring attractor network [57]) as the animal changes heading, thus allowing it to preserve an allocentric reference frame for its heading (and also bearings to objects etc.).

This does not, and need not, imply that each individual knows which way is north, or that different individuals share a common allocentric frame of reference. Instead, it means that individuals use available sensory information to maintain allocentric bearings, including of their own heading [57], with respect to one, or more, stable environmental features. Therefore, while bearings towards other objects in space (e.g., targets) are egocentric in terms of their point of origin—centered on the animal — their bearings are typically encoded in an allocentric (polar) reference frame (see Fig. 1). We stress that this does not imply that the animal has a cognitive map or knows where objects are in absolute allocentric (e.g., Cartesian) space, only that the animal has an allocentric representation of its own heading, and thus also bearings towards represented objects with respect to that heading.

Here, we explore how the egocentric and allocentric representation of space impacts animal movement both in asocial and social contexts. We begin by considering how the internal dynamics of the ring-attractor network give rise to spontaneous patterns of movement in the absence of any sensory input (e.g., during search). This highlights the intrinsic collective neural dynamics (demonstrating the existence of a “critical” transition in neural dynamics), and how intrinsic noise, the strength of neural coupling, as well as whether individuals employ an egocentric or allocentric frame of reference, impact the types of movement exhibited.

Following this, we explore how the neural dynamics, and thus movements, are influenced by simple sensory inputs. Specifically, we consider target-seeking behavior, both in response to static and moving targets in egocentric and allocentric representation. This relates the critical dynamics of the ring-attractor to information processing.

Finally, we consider the emergence of collective motion in such cognitive agents; here the individuals are themselves salient sensory inputs to each other’s ring-attractor network. Thus, the sensory input for each individual becomes much more complex, due to the geometric input to the ring depending both on self-generated motion and the movements of others. In doing so we will demonstrate how collective motion can emerge directly from navigational circuits, and how egocentric and allocentric representations impact such dynamics. Notably, we find that an allocentric representation of space naturally results in the ability for animals to form coherent, mobile groups that exhibit a rich set of patterns, as well as the establishment of long-range order in a manner that is largely independent of density, a property in contrast with some traditional models (such as the Vicsek model [8]). The rich behaviors evident in, what we term “allocentric flocking”, as well as the natural emergence of collective motion from neurobiological principles calls for a shift in perspective, and a new class of models, in the study of collective motion.

## Results

### A minimal ring-attractor model of spatial decision-making

Our model considers an agent equipped with sensory capabilities and decision-making processes governed by a ring attractor neural network of N*_s_* groups of neurons. Following a modelling approach introduced by Hopfield, according to which collective firing activity of neuron groups is modelled by spin variables [40, 68], we assume that the activity of each group of neurons can be represented by a spin variable which can take two states, active (+1) and inactive (*−*1). The state of the spin variables is determined by inputs from other spins as well as by sensory input. Sensory input is represented by excitation induced by the perception of objects in space, with the position on the ring corresponding to the bearing towards the respective object (this can be thought of as each object inducing an external field on the ring). The dynamics of the neural network are governed by a Hamil-tonian, 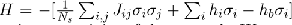, favoring states that minimize energy. Here, J*_ij_* is the synaptic connectivity of the network. We consider a modified cosine-shaped synaptic connectivity (being narrower than a cosine, as supported by previous work [40]). h*_b_* is a background field favoring inactive state, thus representing the degree of global inhibition.

In addition to the input from other neurons, as noted above, each neuron group, i, receives input from the external world, such as a target, in the form of a receptive field, h*_i_*, acting on the neuron. h*_i_* is a function of the angular deviation between the neuron’s receptive field center, α*_i_*, and the target stimulus direction, modeled with a Gaussian response function, with an amplitude of receptive field, h_0_, and standard deviation (receptive field width),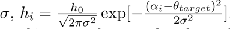.

The agent’s movement is dictated by the activity of its neural network, where the velocity vector is computed based on the activity of the neurons. We consider both whether the resulting direction vector, α*_i_*, is represented in an egocentric, or an allocentric, reference frame (see Fig. 1). See Methods for details.

### Spontaneous neural dynamics

First, we consider the intrinsic internal dynamics (i.e., spontaneous pattern formation on the ring structure). We find that the system exhibits an order-disorder transition as a function of intrinsic (i.e., neural) noise, where the β parameter is the inverse of noise; thus low β corresponds to high noise, and vice versa; see the Supplementary Information S.1.

For values of β below the critical transition point, the system is in the disordered phase, and there exists no correlated activity between adjacent locations on the network. Above the critical point, as β increases (and thus noise decreases), order increases and correlations emerge, but there is no stable (persistent) single bump of activity. For large values of β, however, we enter an ordered phase where adjacent spins (i.e., neural activity) assume similar states and a persistent bump of activity on the ring is observed. For details regarding the degeneracy of the ordered states and the dynamics at the critical point see the Supplementary Information, S.1.).

### From spontaneous neural dynamics to individual motion

How these intrinsic dynamics translate to animal movement depends on whether individuals maintain an egocentric or allocentric representation of space. This difference is not immediately evident for low values of β, such as below, or near, the critical point (Fig. 2**A**). This is because, in this regime, a bump of activity is highly unstable resulting in agents’ motion being slow and highly stochastic (similar to a random walk) for both egocentric and allocentric representations.

**Figure 2:**
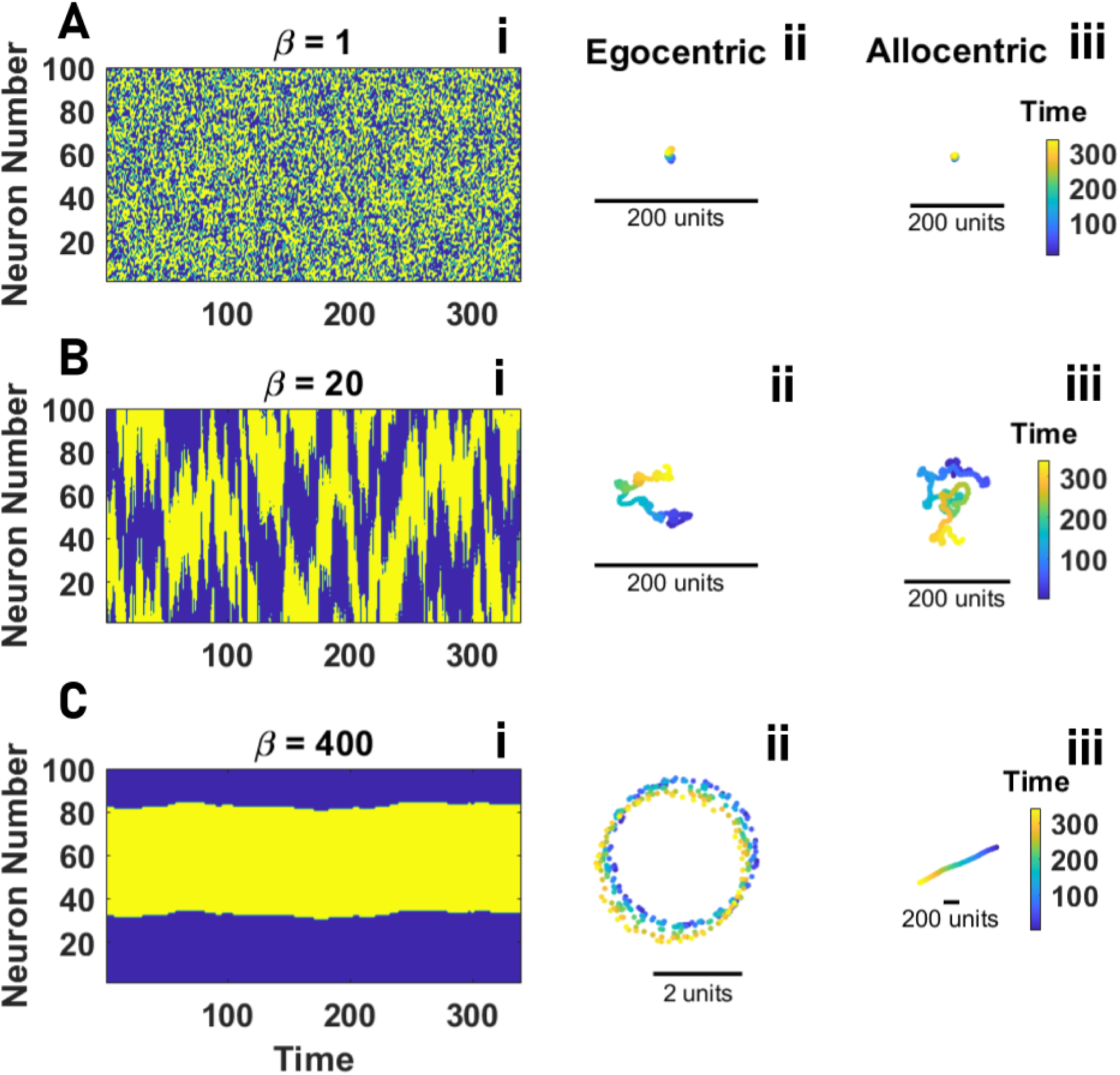
Individual motion. **A** to **C**: The network activity as a function of time (**i**) and the resulting trajectories for egocentric **ii** and allocentric **iii** representation of space, for increasing values of β from the disordered phase (small β, **A**) to the ordered phase (large β, **C**) are shown. In the disordered phase, the agent exhibits a random walk, and no difference between an allocentric and an egocentric representation of space is observed (**A**). As β increases, differences become apparent. An egocentric representation of space results in a more meandering motion (**ii**), and an allocentric representation leads to a more directed motion (**iii**). For larger values of β, corresponding to the highly ordered network activity (**C**), motion patterns with an egocentric representation of space correspond to circular orbits (**ii**), and for an allocentric representation corresponds to a straight line (**iii**). Parameter values: v_0_ = 10, σ = 2π/N*_s_*, h_0_ = 0, h*_b_* = 0, L = 1000, and N*_s_* = 100.

However, as β increases, and the neural activity begins to exhibit a more stable bump (Fig. 2**B****(i)**), different patterns of movement associated with each representation become evident at any spatial scale. For large β, but not too large such that the agent movement is still noisy, with an egocentric representation, the agent tends to often spend long times exploring small regions, with intermittent large jumps (Fig. 2**B****(ii)**). Such a motion is not observed for an allocentric representation of space (Fig. 2**B****(iii)**).

As β increases further still (2**C(i)**), the differences between the agent’s motion with an egocentric and allocentric representation become most evident. For an egocentric representation, agents’ motion tends to an imperfect circular trajectory (Fig. 2**C****(ii)**), whereas for an allocentric representation it is an imperfect directed path (Fig. 2**C****(iii)**). Trajectories approach a perfect circular or directed path, respectively, only for the noiseless infinite β limit. To make this comparison most directly, we illustrate how exactly the same neural dynamics result in very different types of motion: if neural activity is encoded in an egocentric way, a consistent bump position on the ring, α (corresponding to a deviation α of the neural bump from straight ahead), requires the agent to constantly turn with respect to its direction, leading to a circular trajectory with radius R = v_0_/α (if α = 0, however, the agent will move in a straight path in its heading direction). By contrast, if the ring attractor encodes angular information in an allocentric frame of reference, the position of the neural bump is independent of heading, leading to a directed trajectory along an angle α with respect to the agent’s world coordinate system (e.g., an allocentric environmental cue).

### Response to an external sensory input - a ‘target’

Now that we have an understanding of how the internal dynamics result in motion, we can investigate how an external sensory input to the ring-attractor network influences movement. We first consider the simplest case of a single, attractive, static, spatial ‘target’ (below, we extend this to consider response to a mobile target). Congruent with neurological findings [34–37], we assume that such attractive cues induce increased neural activity in the respective region of the ring-attractor.

Even if the target itself is static, we can nonetheless consider this to be an information acquisition problem, since the neural activity on the ring relates both to intrinsic dynamics, and the induced dynamics that are influenced by the agent’s motion, which may change the angular position with respect to the target, and thus, the induced activity on the ring. Thus, even the simplest form of target-seeking is an embodied process, where the motion of the individual may impact the geometric representation of the target, which, in turn, impacts the network activity and thus the resulting individual motion, and so on. This is compounded by the dynamics induced by the motion of the target with respect to the individual in the mobile target case.

We are interested in two aspects of target-seeking; (1) how quickly a target is ‘found ‘ (i.e., how quickly an individual comes into close proximity to a target, following which it may employ a ‘stopping rule’ [69, 70]), and (2) how well an individual can maintain close proximity over time (i..e how well it can track a target). For both egocentric and allocentric representations, individuals move towards the targets. However, the representation employed results in differences between the types of trajectories exhibited. We begin by presenting movement patterns for a fixed target.

### Finding and staying close to a static target

The network dynamics and the trajectory resulting from those dynamics for β values ranging from the disordered phase to the ordered phase, when the agent faces a fixed target, are presented in Fig. 3. For high noise (small β), below the critical point, agents’ movement is predominantly random. However, the external input on the network can induce weak selective movement towards the target. Consequently, agents tend to move slowly toward the target, with speed increasing as a function of increasing β. In this regime, we do not observe differences between egocentric (Fig. 3**A**) and allocentric (Fig. 3**B**) representations since each agent’s movement is predominantly random. See the Supplementary Information for details).

**Figure 3:**
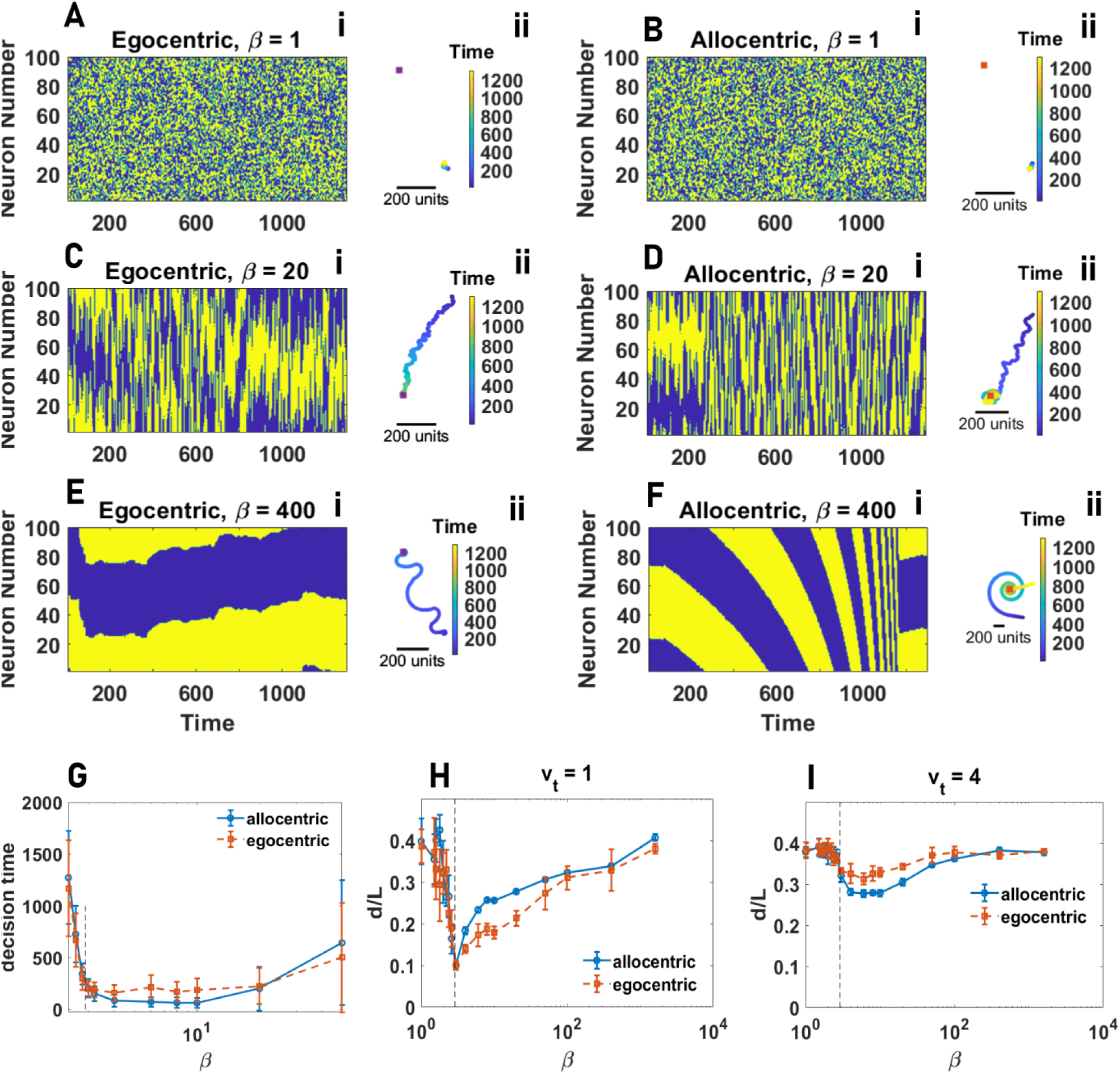
Individual information acquisition. **A** to **F**: The network activity as a function of time (**i**) and the resulting trajectories (**ii**) for egocentric and allocentric representations for increasing values of β from the disordered to the ordered phase is shown. For too small β corresponding to the disordered phase, for both egocentric and allocentric representations of space, the agent only exhibits random and slow movement. Above the order-disorder transition, the agent moves towards the target. For smaller values of β, noise derives transitions between states, which facilitate information acquisition by endowing the agent with flexibility. In the ordered phase, external stimuli result in different network activities for allocentric and egocentric representations of space. With an allocentric representation, external stimuli can lead to the formation of damped traveling waves corresponding to spiral motion toward the target (with more stability for larger values of β). For too large β, the agent occasionally loses interest in the target and moves away. With an egocentric representation, external stimuli help the stability of a bump of activity, allowing the agent to remain stationary once it finds the target. **G**: The agent’s decision-making time in finding a stationary target is plotted as a function of β. Decision-making time is optimized in the ordered phase. An allocentric representation can improve the decision-making speed in the optimal region. **H** and **I**: The average distance of the agent to the target as a function of β for both allocentric and egocentric representations and for different target speeds are plotted. For a stationary or slowly moving target, an egocentric representation is beneficial by allowing the agent to stay stationary once it finds the target. However, for larger target speeds, an allocentric representation improves information acquisition by facilitating the tracking of a fastly moving target. In both cases, the information acquisition optimizes in the ordered phase but is close to the criticality. Parameter values: v_0_ = 10, σ = 2π/N*_s_*, L = 1000, h_0_ = 0.0025, and h*_b_* = 0. In **A** to **F**, N*_s_* = 100, and in **G** to **I**, N*_s_* = 400.

As β exceeds the critical point, we begin to observe clear differences in the trajectories exhibited by egocentric and allocentric agents, with these differences becoming increasingly visible as we move towards the relatively high values of β that characterize the ordered regime. Notably, if employing an egocentric representation, the neural input corresponding to the detection of the target tends to stabilize the bump, and individuals employ a meandering, but relatively direct, path toward the target, and stay confined to a small region when reaching a target (Fig. 3**C** for moderate and 3**E** for large values of β).

If employing an allocentric representation, by contrast, a bump of activity represents a specific bearing in the world. The presence of a target can destabilize this bump. Such a destabilizing effect is not observed for moderate values of β (compare Fig. 3**C** and Fig. 3**D**). For larger values of β, due to the destabilization of the bump, instead of the smooth transitions in the network state that we observed in the absence of an external stimulus, we now find that the network dynamics show sudden transitions between different bumps. This results in a rich set of patterns of motion, such as inward spiraling motion towards the target, corresponding to damped traveling waves of the bump on the ring-attractor network (Fig. 3**F**).

In addition to quantifying the trajectories, we can also ask how individuals may be able to optimize their ability to locate a target. For the fixed target scenario, the agent faces an easy task. In such a simple environment, successful information acquisition can be achieved by simply finding the target and then remaining close to it. We find that target seeking is optimized when the neural network is near the critical point (see S.3). While allocentric and egocentric representations of space perform equally well in such a simple task, if close to criticality, egocentric agents outperform allocentric ones in the ordered phase (see the S.3). This is due to the fact that agents with egocentric representation, once find the target, can stay close to the target by slowing down and/or settling on an attractor with a small radius. On the other hand, an allocentric representation of space can make such a simple task unnecessarily difficult as allocentric agents need to constantly transition between their attractors and update their beliefs about the relative position of the target. This can lead to reduced performance. While finding and remaining close to a target can be of importance in many contexts, in others, successful decision-making may require the agent to only find a target. For example, an animal may consume the target, or the animal may have a stopping rule that they employ once they reach the target [69, 70]. Thus, it is important to address how fast the agent can find a target. To address how allocentric and egocentric perceptions of space affect decision-making time, in Fig. 3**G**, we present the decision time required for the agent to reach a static target. The results indicate that the decision-making time is optimized in the ordered phase. Furthermore, the agent’s decision-making speed is higher (i.e., time taken is lower) with an allocentric perception of space. In the Supplementary Information we show that when higher accuracy in finding the target is required, that is when successful decision-making requires the agent to reach a closer proximity of a target, an egocentric representation of space can become more advantageous for large values of β (S.3).

### Tracking a moving target

We move on to the problem of tracking a moving target. When the target speed is sufficiently small compared to the agent’s average speed, the situation is similar to a fixed target. This is illustrated in Fig. 3**H**, where the average distance of the agent to the target is plotted. However, the situation changes in a fast-changing environment, where the agent needs to track a target with an appreciable speed relative to the agent’s average speed. In Fig. 3**I**, we present the distance of the agent to a moving target with a high speed. While for a slowly moving target, egocentric agents can outperform allocentric ones, with a high target speed, the existence of an allocentric representation provides a benefit because, when employing an egocentric representation, the agent’s movement can lead to dramatic fictitious changes in the external world, brought about by the shifting agent’s position (we note that, this is not a problem when seeking a fixed target, because in such cases, the agent can use a simple strategy of standing still once finding the target). Accounting for these changes requires large changes in the network activity to constantly encode a mobile target’s position. On the other hand, the environmental change resulting from the agent’s movement is not dramatic from an allocentric perspective. Consequently, allocentric agents can easily modify their movement trajectory by small shifts in their bumps of activity.

We also find that, while higher values of β decreases time to reach the target (which can be considered decision-making speed), the ability of the agent to stay in close proximity to the target (which can be considered decision-making accuracy) is higher for lower values of β. This trade-off is optimally solved not at the critical point, but in the ordered phase. This is due to the fact that following a moving target requires a more coherent movement of the agent, which can only be achieved in the ordered phase. Furthermore, the distance of the optimal decision-making region to criticality increases as the target’s speed increases.

### Collective motion

Above, we demonstrated that even for simple sensory inputs, having either an egocentric or allocentric neural representation in the ring attractor network can greatly impact movement. Now we consider the far more complex sensory environment experienced by individuals in social groups and ask how egocentric and allocentric representations of bearings towards others impact collective movement. Here, the individuals themselves become static, or mobile, targets from the perspective of others. Thus, the neural ring attractor dynamics of each individual both influences, and is influenced by, that of others, as a recursive (recurrent) feedback loop.

The model of collective movement straightforwardly results from the individual movement model by having a population of N agents, each of which is a target for others, with an amplitude of the receptive field, h*^s^*. Collective movement can thus be studied using a single parameter, a social attraction parameter, h*^s^*, which parametrizes the strength of social attraction (see Methods). As the control parameter of the model, we consider the total social attraction, h*^t^*, defined as the social attraction of an individual towards another individual, h*^s^*, times the population size, N. As we will see, normalising social attraction acting on an individual due to each other individual, h*^s^*, by the population size results in a similar phase diagram for different population sizes. This is implemented by taking the control parameter of the model to be h*^t^* .

In the main text, we focus on the global order and local order of the system. Global order (GO) is calculated as the sum of the normalised velocity vectors of all individuals (equivalent to the order parameter of the Vicsek model [8], and the alignment/polarisation of the system as employed in collective behavior studies [5]). Local order (LO) is the average normalised velocity, not of the whole system, but of each individual’s local topological neighborhood. This allows us to differentiate, for example, between disordered dynamics at all scales, such as when there is disorder, and thus low local or global alignment, and states where there is low global order, but high local order, such as when populations are composed of multiple small, coherently-moving groups, but each tends to move in a different direction. See Methods for details and Supplementary Information, S.4, S.5, and S.6 for the supplementary analysis (e.g., using measures of distance between agents).

### Egocentric representation of space

We begin by considering the emergence of collective motion for individuals that employ an egocentric representation. In Figs. 4**A** and 4**B**, we plot the global and local order parameters in the plane defined by β and h*^t^* . As the strength of social attraction among individuals increases, we see that populations cannot achieve global order (Fig. 4**A**), and thus large-scale collective motion is never observed. However, we find that local order tends to be moderate to high (Fig. 4**B**). Thus, agents’ direction of travel is similar to their close neighbors, but this emergent alignment is highly localised. At the scale of the population, increasing social attraction results in the formation of aggregation (but not collective motion), where agents coalesce in a dense group with low mean nearest neighbor and all pair distance (see S.5).

**Figure 4:**
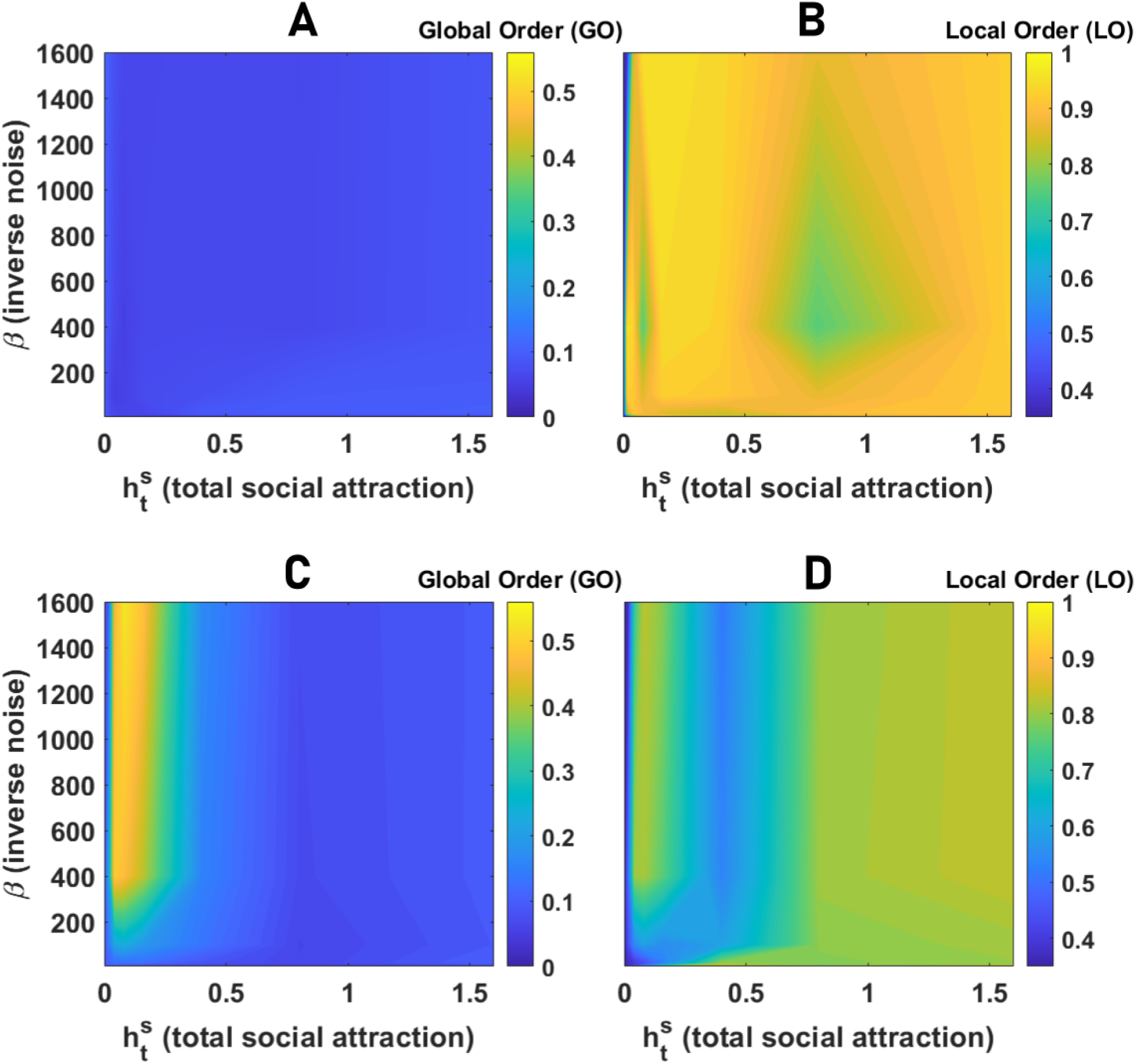
Collective behavior of agents with egocentric and allocentric representation of space. **A** and **B**: Global order (GO in **A**), defined as the angular order parameter (AOP), and Local Order (LO in **B**), defined as the topological vectorial order parameter (VOP), in groups of 80 agents with an egocentric representation of space are color plotted as a function of the network inverse temperature, β, and total social attraction, h*^s^*. For a too-small social attraction, the agents move independently. As the social attraction increases, local order increases but not global order, indicating the onset of an aggregation phase where agents aggregate in a stationary dense group. **C** and **D**: The global (**C**) and local (**D**) order in groups of 80 agents with allocentric representation of space as a function of the network inverse temperature, β, and total social attraction, h*^s^*, are color plotted. The system shows three distinct phases: disordered motion for small h*^t^*, collective motion with high local and global order, and aggregation phase with low global but high local order. Local order is minimized close to the phase transition between collective motion and aggregation due to the strong fission-fusion dynamics leading to explosive movement of the densely packed group. Parameter values: N*_s_* = 100, v_0_ = 10, σ = 2π/N*_s_*, h*_b_* = 0, N = 80, and L = 1000.

While the lack of global order is manifest in the insensitivity of GO to variation of social attraction (Fig. 5**A**), LO shows bimodality close to the phase transition (Fig. 5**B**). This shows that this phase transition is discontinuous (see S.5 for details). A high value of local order in the ordered phase indicates that agents’ heading points in a similar direction compared to their neighbors. However, this local order does not induce collective motion. Instead, agents form a compact group. A snapshot of the agents in this phase is presented in Fig. 5**C**. For smaller values of β, presented in Fig. 5**D**, a similar situation is observed, however, in this case, noise in individual agents’ movement derives a random walk-like motion within the cohesive aggregation of agents.

**Figure 5:**
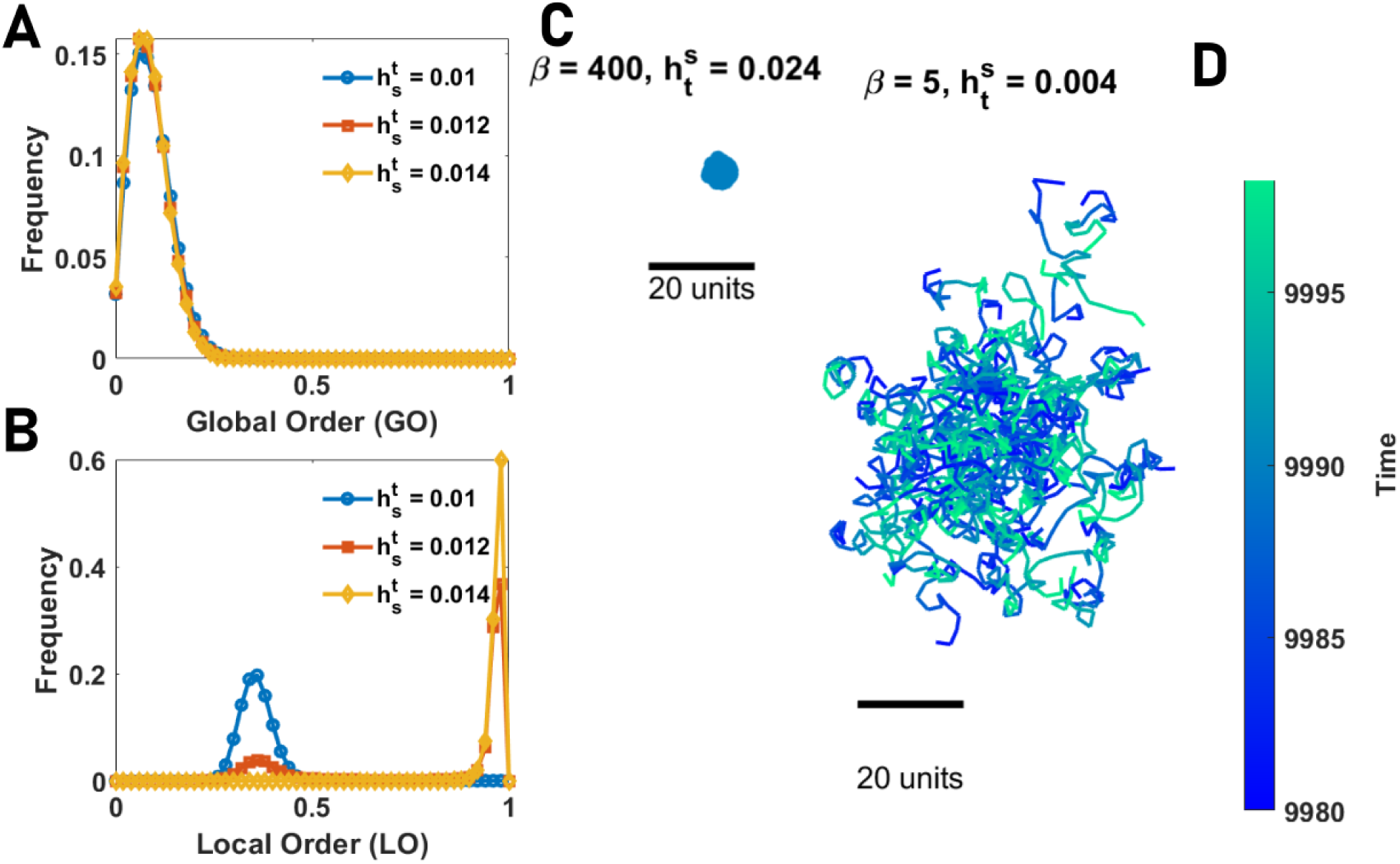
Phase transitions in collectives with an egocentric perception of space. **A** and **B**: The distribution of global order (GO) and local order (LO) in groups of various sizes of agents with egocentric representation of space. While GO takes a small value expected in the disordered phase and does not show sensitivity to social attraction, LO shows bimodality close to the order-disorder transition, indicating a discontinuous transition. **C** and **D**: Snapshots of the collective behavior in the ordered phase are shown. For large β (low network noise), in the ordered phase, agents form an almost stationary circular pack of densely aggregated agents. For smaller β the pack’s radius increases and agents perform a random-walk-like movement within the pack. Local order is high in both cases. Parameter values: N*_s_* = 100, v_0_ = 10, σ = 2π/N*_s_*, h*_b_* = 0, β = 400, N = 80, and L = 1000.

### Allocentric representation of space

The situation changes dramatically if agents possess an allocentric representation. Now three distinct regimes are observed. This can be seen in Fig. 4**C** and 4**D**, where the GO and LO as a function of β and h*_s_* are plotted. As social attraction increases, the system shows a phase transition from a disordered phase to a phase where both local and global order are high, indicating the emergence of large-scale collective motion out of agents’ inclination to stay with the group. Further increasing h*_s_*, yet another phase transition from the collective motion phase to an aggregation phase where agents form a relatively immobile aggregation with high LO and low GO is observed.

For relatively high neural noise (low β), large-scale collective motion is never observed, as we only see a cross-over from the disordered phase to the aggregation phase where agents form a compact pack in which local order is observed, but no collective motion emerges (Fig. 4**C**). This is because, in our model, collective motion is a collective information acquisition problem and emerges due to agents coming to a consensus regarding their direction of travel. Thus, coherent large-scale collective motion requires agents to be able to keep a spatially-consistent bump of neural activity over time, which can only occur if neural dynamics are not too noisy (i.e., are in the ordered phase, Fig. 2**F**). We term this “allocentric flocking”.

### Allocentric flocking and population (system) size

By studying the statistical properties of the different regimes of allocentric flocking, we find that the motion patterns observed are sensitive to system size.

For very small system sizes, such as N = 10, the order-disorder transition (the transition from disordered motion to collective motion) exhibits bistability. This can be seen in Fig. 6**A**, where the distribution of global order close to the disorder-order transition shows two peaks, one at low and the other at high, GO (representing the disordered and ordered phase, respectively). This indicates intermittency between disordered motion and ordered motion. We find that agents show a wide range of collective motion patterns including swirling, sudden expansions (similar to ’flash expansions’ exhibited by animal groups [71–73]), fission-fusion dynamics, as well as coherent, directed motion. Fig. 6**E** presents a snapshot of motion patterns during swirling when the group rotates around a common origin, and Fig. 6**F** for a swirling resulting in a coil-shaped trajectory. See the Supplementary Videos 1 and 2. In this regime, global order is low, but local order is high and the distance between agents exhibits strong fluctuations over time. This can be seen in the blue line in Figs. 6**C** and 6**D**, where GO and mean distance between all pairs for different values of h*^t^* are plotted.

**Figure 6:**
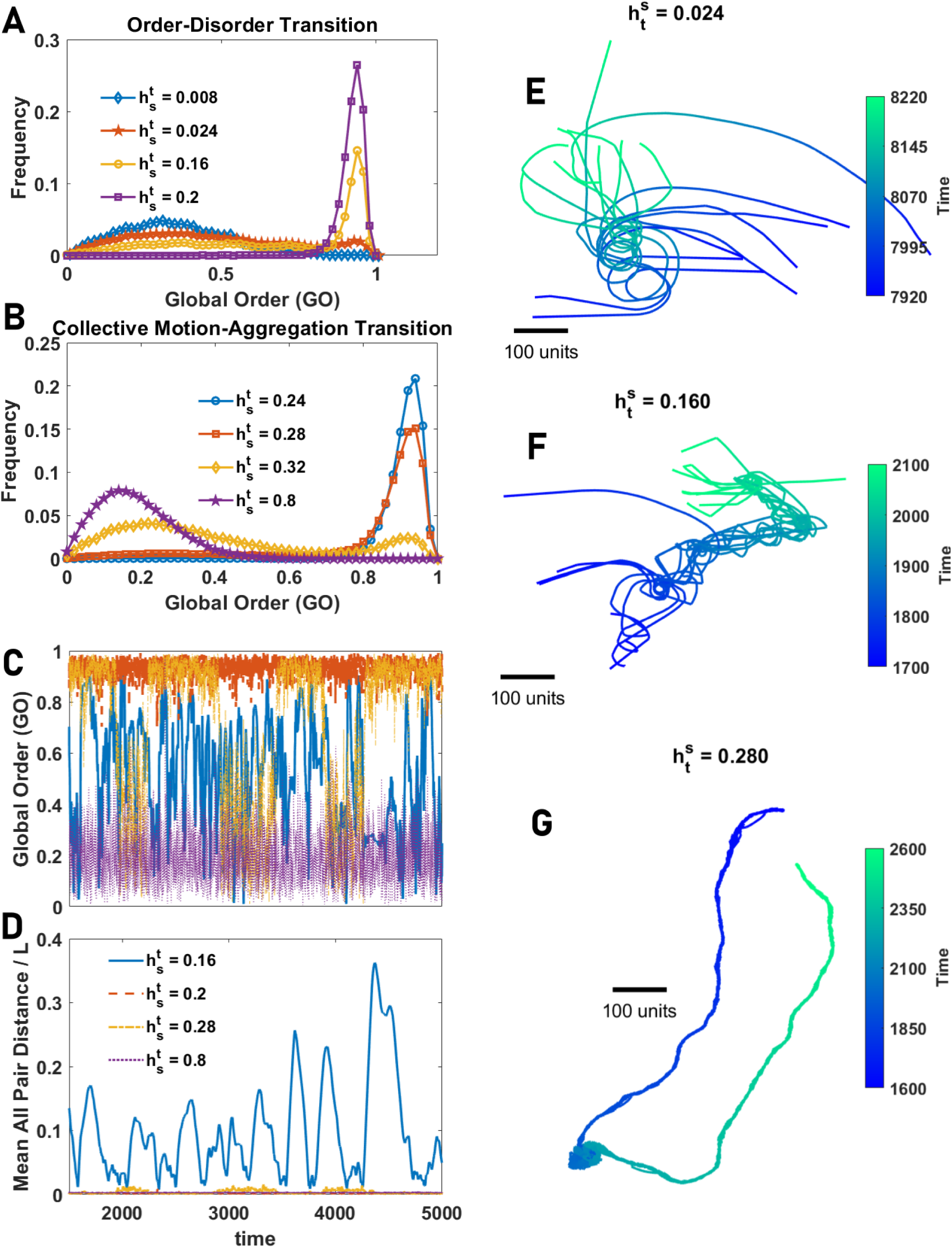
Collective behavior in small groups of agents with an allocentric representation of space. **A** and **B**: The distribution of GO and LO in groups of various sizes of agents with allocentric rep- resentation of space close to the order-disorder transition (**A**) and close to the collective motion- aggregation phase transition (**B**). In both cases, the distribution shows bimodality signaling a discon- tinuous pseudo-phase transition in small system sizes. **C** and **D**: GO and mean distance between all pairs normalised by space size, L, as a function of time for different values of total social attraction are plotted. For small social attraction, the system shows intermittency between high and low order, and for larger social attraction intermittency between ordered motion and aggregation is observed. **E** to **G**: Example snapshots of motion patterns from the Supplementary Videos for some values of *^s^* shown in **A** and **B**. In the collective motion phase, the system shows a rich set of motion patterns, including swirling in circular orbits (**E**), or in coil-shaped orbits (**F**), fission-fusion dynamics and intermittency between highly ordered motion and aggregation (**G**). Parameter values: N*_s_* = 100, v_0_ = 10, σ = 2π/N*_s_*, h*_b_* = 0, β = 400, N = 10, and L = 1000.

In Fig. 6**G** we present a snapshot of the motion patterns for directed motion in relatively small groups (N = 10, see Supplementary Video 3). In this example, agents form a coherent, mobile group (as exemplified in Figs. 6**C** and 6**D**, red line). This trajectory also shows an example of intermittency between such directed motion and the formation of a stationary aggregation (where global order transiently decreases (orange line in Figs. 6**C** and 6**D**), resulting in a (probabilistically- likely) reorientation once the group transitions back to coherent motion. This aspect of collective behavior is reflected in the bimodal distribution of local and global order parameters, as shown in Fig. 6**B** (see the S.4 for details).

As population size increases, however, the situation changes. While transitions in collective state appear discontinuously in small system sizes, they become more continuous for larger system sizes. This is shown in Fig. 7**A** and 7**B** for the disordered-collective motion, and collective motion to aggregation phase transitions, respectively (Here N = 320. See S.5 for other system sizes). In larger populations, intermittency between different motion patterns is more frequent. Furthermore, due to the prominent fission-fusion dynamics, the population is more likely to be decomposed into different groups exhibiting different collective motion patterns, such as collective motion, swirling, explosive movement, or sudden direction changes. This results in global order being a less effective means of characterising the collective behaviours exhibited by agents. Examples of some motion patterns, including collective motion and fission-fussion dynamics are presented in Figs. 7**E** and 7**F**. See the Supplementary Videos 4, 5, 6 and 7. Notably, to reach different motion patterns, it is not necessary to tune the parameters of the model; Rather diverse motion patterns occur for the same parameter values and are exhibited by the same population of individuals over time.

**Figure 7:**
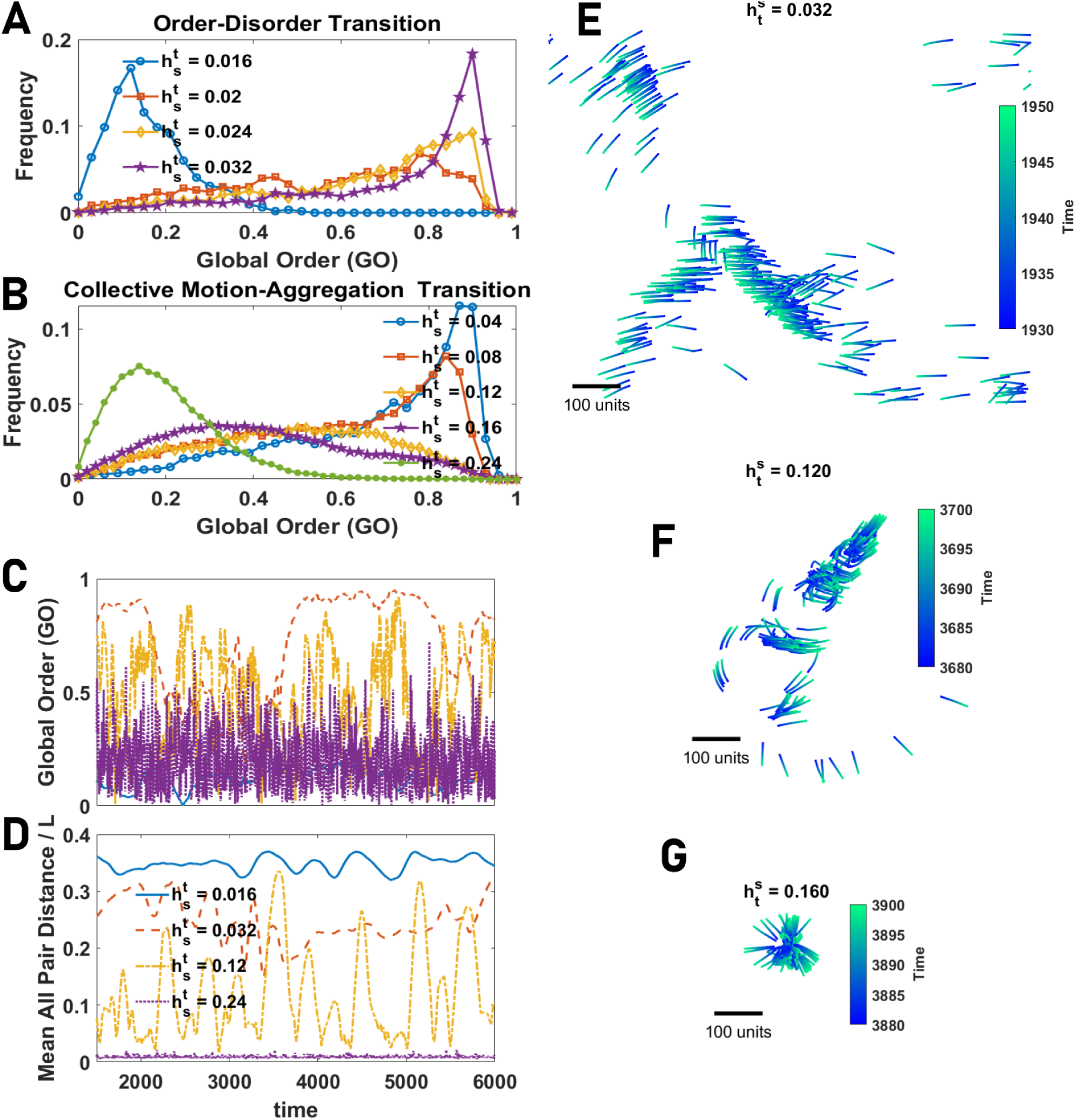
Collective behavior in large groups of agents with allocentric representation of space. **A** and **B**: The distribution of AOP and normalised topological VOP in groups of various sizes of agents with an allocentric representation of space close to the order-disorder transition (**A**) and close to the collective motion-aggregation phase transition (**B**). Both phase transitions tend to a continuous phase transition, indicated by large fluctuations and broad distribution of the order parameter, as group size increases. **C** and **D**: Global order (GO) and mean distance between all pairs normalised by space size, L, as a function of time for different values of total social attraction are plotted. The system shows intermittency between high and low order resulting from transitions between different motion patterns and strong fission-fusion dynamics. **E** to **G**: Example snapshots of motion patterns from the Supplementary Videos for some values of h*^s^* shown in **A** and **B**. In the collective motion phase, the system shows a rich set of motion patterns, including flocking (**E**), sudden direction change and fission-fusion dynamic (**F**), and intermittency between highly ordered motion and aggregation leading to explosive movement with low local order (**G**). Parameter values: N*_s_* = 100, v_0_ = 10, h*_b_* = 0, σ = 2π/N*_s_*, β = 400, N = 320, and L = 1000.

By increasing the social attraction the system shows a continuous phase transition to an aggregation phase where the population forms a dense group of agents lacking directed motion. However, near the transition, frequent explosive and implosive movements of the population are observed (this results in a decline in local order in the phase transition region, as observed in Fig. 8**A**). A snapshot of the motion pattern in this phase is plotted in Fig. 7**G**. See the Supplementary Video 8.

**Figure 8:**
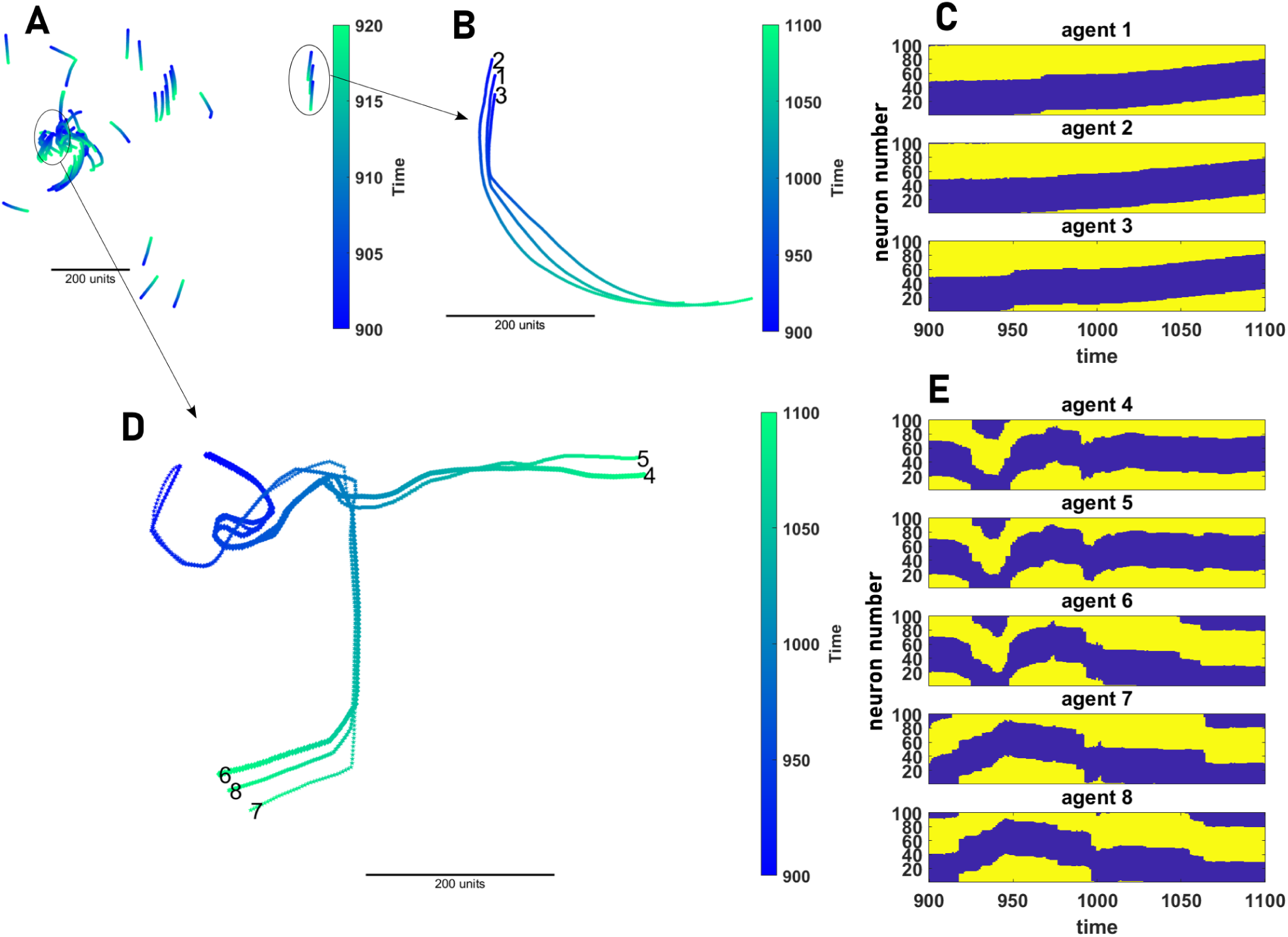
Cognitive representation of collective motion. **A**: A snapshot of the collective motion in a population of 80 agents is shown. The population can be decomposed into subgroups of synchronized agents. **B** and **C**: The motion pattern (**A**) and the neural activity (**B**) of a subgroups of 3 coherently moving agents among the 80 agents presented in **A** are shown. The coordinated movement of the agents results from the synchronization of their neural dynamics. **D** and **E**: An example of fission- fusion and leader-follower dynamics is shown. In the beginning, agents 4 to 6 are synchronized and move together, and agents 7 and 8 move together. When these two groups come into close proximity, at around timestep 1000, agent 6 “changes its mind” and joins agents 6 and 7. Consequently, its neural activity becomes synchronized with agents 7 and 8. Around time 1050, a sudden direction change by agent 6 drives a sudden direction change in agent 8, followed by a similar behavior of agent 7. Parameter values: N*_s_* = 100, v_0_ = 10, σ = 2π/N*_s_*, h*_b_* = 0, β = 400, N = 80, and L = 1000.

### Cognitive representation during collective movement

In our model collective motion results from simple feedbacks between the ring-attractor networks employed for spatial navigation by animals. Key to the patterns observed in the collective context, such as sudden and coordinated changes in direction of mobile groups, is the synchronization of the neural dynamics of the agents. In Fig. 8, we present a snapshot of collective motion in a population of 80 agents. The population can be decomposed into subgroups of coherently moving agents whose neural dynamics exhibit synchronization. Evaluating the spatiotemporal dynamics of the neural representation in the “brain” of each agent, we see there exist emergent leader-follower dynamics, as evident by direction changes of an individual (change in the position of the neural activity bump on their ring attractor) being followed by similar changes in others (Figs. 8**B** to 8**C**) or fission-fusion dynamics, whereby an individual (or subgroups) desynchronise (resulting in fission) or synchronise (resulting in fusion) with others (Figs. 8**D** and 8**E**). See the S.7 for more details.

### Parameter dependence

In Figs. 9**A** and 9**B** we look at density dependence by changing the size of the arena. Here, we compare GO and LO for N = 80 agents in a square arena with periodic boundaries of linear dimensions, L = 1000, and L = 10000 are compared. We observe that, in contrast to classical models based only on local alignment [8], density does not affect the phase transitions. Rather, similar phase transitions at similar values of social attraction are observed at both low and high population densities, indicating a lack of density-dependent phase transitions. See the S.6 for details.

**Figure 9:**
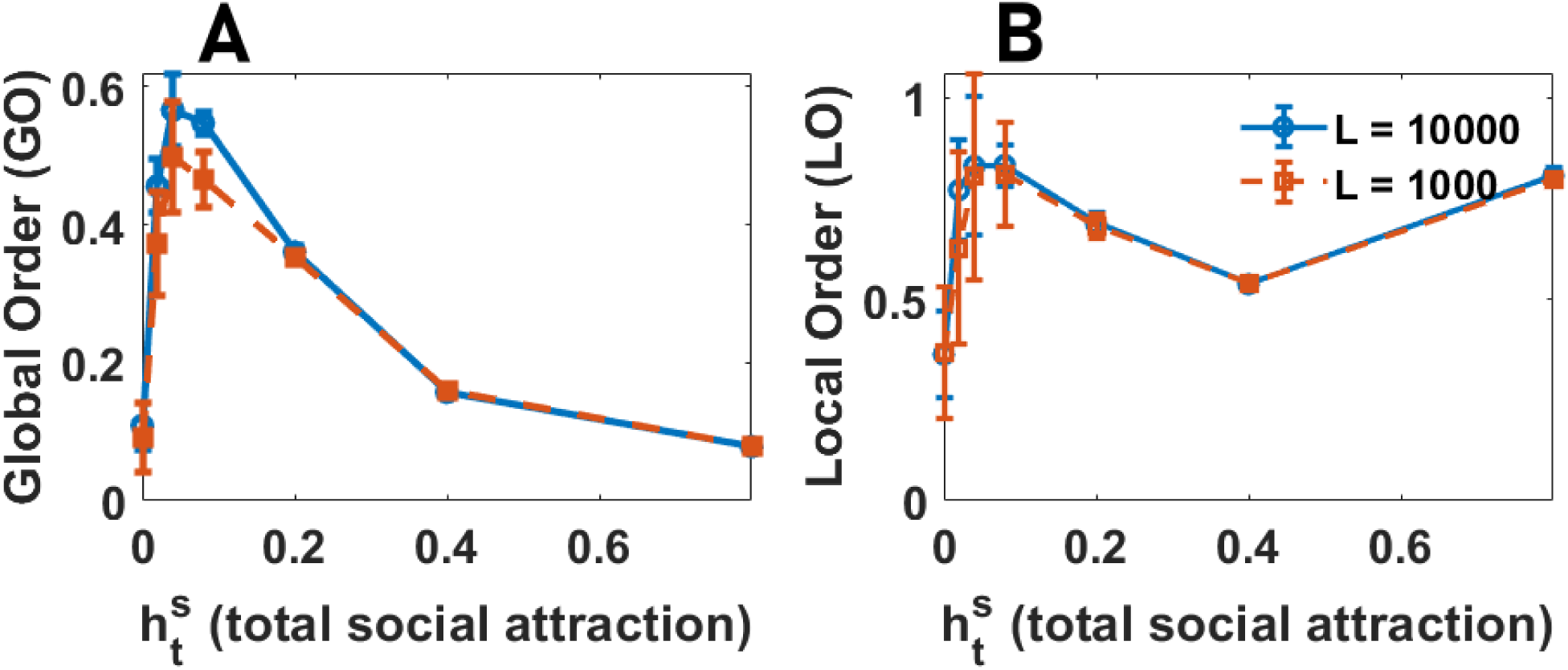
Lack of density dependence phase transition in agents with allocentric representation of space. Global order (GO in **A**) and local order (LO in **B**) as a function of total social attraction for groups of 80 agents with allocentric representation of space in a periodic space with linear size L for two different values of L, leading to different mean density, are plotted. The same phases and phase transitions are observed for different densities, indicating a lack of density dependence. Global order increases in large system sizes due to less frequent mixing of coherently moving front in free space. Parameter values: N*_s_* = 100, v_0_ = 10, σ = 2π/N*_s_*, h*_b_* = 0, β = 400, N = 80.

In the Supplementary Information we confirm the phenomenology of the model holds for other parameter values, such as the complexity of the agent (the number of spins), speed constant, v_0_, and the width of the receptive field of the agents (S.6). Besides, we show that our findings are robust when short-range repulsion is introduced to the model (S.8), or when social attraction decays with distance (S.9).

## Discussion

Here, we have shown that a rich suite of collective behaviours, including the formation of coherent, mobile groups, emerges naturally from the types of neural circuits—ring attractor neural networks— employed by animals during spatial navigation. By contrast to classical models of collective motion, that use hypothetical rule-based interactions, such as repulsion, alignment and attraction, our modelling framework is grounded in cognitive principles of spatial information processing in the (invertebrate and vertebrate) brain. We show that collective motion can emerge directly from navigational circuits, without requiring explicit alignment, or additional rules of interaction — if individuals employ an allocentric (but not an egocentric) representation of space. While not previously considered in the study of collective behaviour, this spatial representation is known to be ubiquitous, employed by fruit flies [57], and humans [58], alike. Following the historical use of the term ’flocking’ to broadly describe the collective dynamics of diverse systems—whether physical particles, animals, or robots [5]—we term this mechanism ”allocentric flocking”.

Allocentric flocking results from interactions among cognitive agents with an allocentric perception of space, where individuals themselves act as sensory inputs to each other’s ring attractor networks. While at an individual level, we find that an allocentric representation of space can be beneficial for effective target-seeking in a rapidly changing environment, it is shown to be essential to achieve coherent collective motion. This provides a contrasting, but empirically-grounded, explanation as to how collective behavior may arise in many animal species, thus shedding light not only on the mechanistic basis of collective motion, but also how it may simply, and thus readily, evolve from an asocial ancestral state.

The introduction of cognitive processes into models of collective behavior demonstrates that local alignment, a common feature attributed to many collective systems, may be an emergent rather than an intrinsic property, a view supported by data from multiple species that exhibit collective motion [54, 55]. It arises in our model as a form of consensus dynamic (not dissimilar, conceptually, to models of collective information acquisition [74]) for agents who have an allocentric representation of bearings (their own heading direction, and the bearings towards others, are within a world-centred frame). We show that, by contrast, if individuals exhibit an egocentric representation (whereby bearings are body-centred, but directional bearings are only encoded with reference to the present heading), social attraction can only result in the formation of relatively immobile aggregations. Here, the additive nature of attraction is analogous to “gravitational collapse” [75].

Our results suggest that the known allocentric polar coordinate system employed in the brain by diverse species, from insects [57] to mammals [58], can allow collective motion to arise from, and be regulated in, navigational circuits. This parsimonious framework demonstrates how easily collective behavior can emerge from known neurobiological principles. It can also readily be modified to be adapted to specific systems to incorporate further features, such as individual and collective learning, or to address different ecological questions, such as collective sensing, navigation and decision-making. By introducing allocentric flocking as a general mechanism for the emergence of collective behavior, we hope to encourage further research into the feedback loop between neural dynamics and organismal collective behaviors.

## Methods

### The Model

We consider cognitive agents capable of sensing and decision-making. Each agent’s decisions are governed by a ring attractor neural network with N*_s_* neuron groups and endowed with a ring structure.

Neuron groups are modelled as spin variables, long used to model neural systems [68, 76], and can take two states, active, +1_L_, and inactive, *−*1. The activity of each neuron, i, is determined by an We take the synaptic connectivity of the network, J*_i,j_*, to be a modified cosine function, as follows:

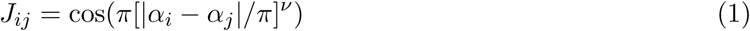

This implies that neurons in the network have periodic connectivity and endow the network with a ring structure. With ν = 1, positive and negative synapses are found in roughly equal numbers, and for ν < 1, the network connectivity is locally more excitatory and globally more inhibitory, which requires more inhibitory synapses to exist in the system.

The external field is determined based on the sensory input the neuron groups receive. Each neuron group, i, has a receptive field centered around the angle α*_i_*. Without loss of generality, we take α*_i_* = 2πi/N*_s_* (the angle α*_i_* is measured with respect to the positive x *−* axis). A neuron group responds to external stimuli based on the angular deviation of the stimuli with its receptive field center:

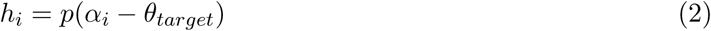

Where θ*_target_* is the angular position of a target (external stimulus) with respect to the agent. p can be thought of as the response function of a sensory neuron sensitive to direction α*_i_*. We will work with a Gaussian response function given by the following equation:

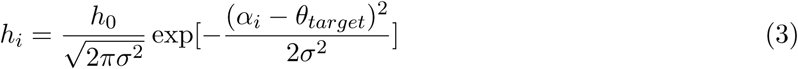

We assume the network dynamic is governed by a Hamiltonian, as follows:

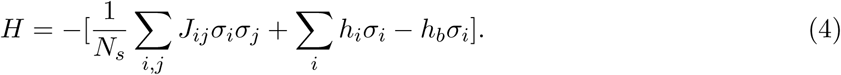

Here, h*_b_* is a constant term that promotes inhibition of the network activity. Assuming neurons favor a state with the lowest energy, this Hamiltonian implies that each neuron group tends to assume a state favored by its input. We use the Glauber dynamic to simulate the network’s dynamics [76]. At each step, a neuron is chosen at random, and the energy difference resulting from updating the neuron’s state is calculated. The neuron’s state is flipped with certainty if the energy difference becomes negative, and it is flipped with probability exp(*−*βΔH) if the energy difference is positive. We repeat the Glauber dynamics for T_0_N*_s_* steps for the network to equilibrate. After this, we update the agents’ position according to the equilibrium activities of the neurons. In this stage, the agent moves by a speed v⃗ determined by the activity of its neural network according to the following equation:

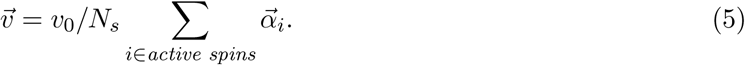

Where, α⃗*_i_*, is a vector pointing toward direction α*_i_*.

We consider both egocentric and allocentric representation of space. While the neural mechanisms using which the brain forms an allocentric representation of space are not fully clear and can vary across organisms, the presence of an allocentric representation of space seems to be universal even in simple organisms [57–62]. This observation points to one of the limitations of previous works on attractor neural networks which needs to be removed [40, 52, 53]. To do so, we do not model the way in which the brain may form an allocentric representation of space. Rather, we take the direction vector, α⃗*_i_*, to be an allocentric direction calculated in a reference frame independent of the agent’s head direction. On the other hand, with an egocentric reference frame, the reference frame is attached to the individual, and thus, rotates as the individual’s head direction changes.

The extension of the model of individual movement and information acquisition to a model of collective movement is rather straightforward and only requires a change of perspective: it is enough to allow several such agents to perceive each other as possible targets and interact. We consider three variants of such a model of collective motion, based on the regulation of social interaction.

In the baseline model, we consider the simplest case, where each agent is a target to other agents, with a magnitude of external field, h*^s^*. We usually report total social attraction, defined as 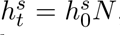 We also consider two variants of this baseline model. In the model with short-range repulsion, the amplitude of the receptive field is a step function of the distance of the focal agent to the target. Below a collision radius, the amplitude of the external field is taken to be a negative value with magnitude, ensuring conspecifics act as a repelling stimulus, rather than an attracting one. Taking the magnitude of social repulsion large enough ensures agents avoid a collision. Above the collision radius, the external field is taken to be positive, ensuring conspecifics are attracting stimuli.

In the second variant of the model, we study the distance dependence of social attraction. In this variant, the amplitude of the receptive field is taken to decay with the distance between the focal agent and its target, d, according to an exponential, h*^s^* exp(*−*d/ζL), where L is the linear size of the space. With this choice, for d < ζl, the exponential term is approximately a constant and equal to 1. ζL is thus, the characteristic length of social attraction, above which the strength of social attraction decays exponentially fast.

### Statistics and Reproducibility

#### Simulations

The base parameter values used for the individual motion patterns are as follows N*_s_* = 100, v_0_ = 10, σ = 2π/N*_s_* (unless otherwise specified). All the simulations are performed in a space with periodic boundaries. Unless otherwise stated, the linear size of the space is equal to L = 1000. For collective movement, agents “see” each other, such that each agent sees each other agents with an amplitude of external field equal to h*^s^*. We report the total external field defined as h*^s^* = h*^s^*N, where N is the population size. In Figs. 3**G**, 3**H**, and 3**I** N*_s_* = 400 and other parameter values remain the same. The averages and error bars in Figs. 3**H** and 3**I** are calculated based on the stationary state of a sample of 5 run for 10000 timesteps. The target’s speed along the x and y axis obeys a random walk with speed v*_t_*, shown on the panels. In 3**G**, we have used 80 simulations, and the simulations stop when the agent reaches a close proximity of the target (5 units). Here, the target is stationary. The use of a larger sample is due to the fact that in such a decision-making speed task, it is not possible to rely on long-time stationary trajectories to provide stronger statistics. The amplitude of the external field in all the cases is equal to h_0_ = 0.0025. Error bars represent the standard deviation over the sample. Large error bars for large values of β in Fig. 3**G** are because the agent’s decision-making accuracy decreases for too large values of β, while its speed increases. Thus, while in some trials the agent reaches the target rapidly by moving directly toward the target, in other trials it starts by moving in the wrong direction.

In Fig. 4, a sample of 3 simulations run for 10000 timesteps in a population of N = 80 agents moving in a space with periodic boundaries and linear size L = 1000 is used. In Fig. 5, simulations are performed for 15000 timesteps. A sample of 10 simulations is used to calculate the distribution. The distributions are calculated based on the last 10000 time steps of the simulations to ensure stationarity. Here, N = 80 and L = 1000.

In Fig. 6 and Fig. 7, simulations are performed for 15000 timesteps. A sample of 5 simulations is used to calculate the distribution. The distributions are calculated based on the last 13500 time steps of the simulations to ensure stationarity. Here, L = 1000, and N = 10 and N = 320 respectively.

In Fig. 9**A** and 9**B** a sample of 5 simulations is used. For L = 1000 simulations are performed for 10000 time steps and for L = 10000, the simulation time has increased to 40000 to increases statistics. Errorbars are standard deviation. Averages and errorbars are calculated discarding the first 5000 timesteps.

#### Measures of collective movement

In this section, we define the measures used to analyze simulation data in our study. Further analysis is performed in the Supplementary Information and a more complete list of the measures used in the study is provided in the Supplementary Information (S. 10).

*•* Global Order: As a measure of global order, we have used the angular order parameter. This is simply calculated as the sum of the normalised velocity vectors of the individuals. To do so, we have normalised direction vectors to 1 and then summed over all individuals’ normalised vectors. Values close to 1 indicate strong alignment and values close to 0 indicate weak alignment. We note that, in practice, the minimum of this quantity approaches zero only in the limit of infinite population size.
*•* Local Order: As a measure of local order, we have used the topological vectorial order parameter. This is a measure of the average direction of the velocity vectors of individuals within a local neighborhood defined based on topological distance, LO = *_i k_* _nearest neighbors_ v⃗*_i_*/k, where the summation is over k nearest neighbors of the focal individual. We have set k = 5. The results are valid for other reasonable choices. A high value indicates strong alignment (coordinated movement in a common direction), while a low value indicates weak alignment.
*•* VOP and geometric VOP: This is similar to topological VOP, however, it is calculated in a local neighborhood based on geometric distance (geometric VOP) or the entire group (VOP). In the case of Geometric VOP, all the individuals closer than a distance R are considered. An average over all the individuals is performed. These measures can take values up to the total speed observed in a neighborhood (or the total population). In all the plots we have taken R = L/100.
*•* Mean distance between all pairs: This is the average distance between all the pairs in the group, *_i,j_* x*_i,j_*/(N(N *−* 1)), where x*_i,j_* is the distance between individuals i and j, and N is the number of

agents in the group.

### Supplementary Videos

Parameter values used in the Supplementary Videos are as follows: N*_s_* = 100, v_0_ = 10, σ = 2π/N*_s_*, β = 400, and L = 1000.

Supplementary Videos SV.1 to SV.6 present the dynamics of collective motion over time. Snapshots of the Videos are presented in Fig. 6 and 7. In SV. 1 to SV. 3 N = 10 agents interact and in SV. 4 to SV. 6 N = 320 agents are considered. Total social attraction in these videos is equal to h*^s^* = 0.024, h*^s^* = 0.16, and h*^s^* = 0.28, respectively. These videos correspond to snapshots presented in Fig. 6**E** to 6**G**, respectively. In Both SV.1 and SV.2 a variety of motion patterns including intermittent swirling, sudden direction change, and fission-fusion dynamics can be observed. SV.3 is chosen close to the phase transition between the collective motion-aggregation phase, and the intermittency between these two modes of motion can be observed.

SV.4 to SV.8 show examples of collective motion in large groups. The total social attraction is set equal to h*^s^* = 0.02, h*^s^* = 0.032, h*^s^* = 0.08, h*^s^* = 0.12, and h*^s^* = 0.16, respectively. SV.5, SV.7, and SV.8 correspond to snapshots in, Fig. 7**E** to 7**G**, respectively. Both SV. 4 and SV. 5 show strong collective motion. However, intermittency and coexistence of different modes of motion, such as fission-fusion dynamics, swirling, startling, and sudden direction changes can be observed. Similarly in SV. 6 and SV. 7 a variety of collective motion can be observed. Explosive and implosive motion of the group leads to highly coordinated state changes between different motion patterns. The explosive and implosive motion is stronger in SV. 8 chosen at the transition between collective motion-aggregation.

## Acknowledgement

The authors acknowledge funding from Deutsche Forschungsgemeinschaft (DFG, German Research Foundation) under Germany’s Excellence Strategy - EXC 2117-422037984, the Deutsche Forschungsgemeinschaft Gottfried Wilhelm Leibniz Prize 2022 584/22 (I.D.C.), the Max Planck Society, the European Union’s Horizon 110 2020 Research and Innovation Programme under the Marie Skl-odowska- Curie Grant agreement no. 860949, the Struktur- und Innovations fonds für die Forschung of the State of Baden-Württemberg, the PathFinder European Innovation Council Work Programme no. 101098722, the Office of Naval Research Grant N0001419-1-2556.

## Author Contribution Statement

M.S. and I.D.C. designed the research, M.S. performed the research, and M.S. and I.D.C. wrote the paper.

## References

[1] Goldstone, R.L. and Janssen, M.A., 2005. Computational models of collective behavior. Trends in cognitive sciences, 9(9), pp.424–430.

[2] Ouellette, N.T. and Gordon, D.M., 2021. Goals and limitations of modeling collective behavior in biological systems. Frontiers in Physics, 9, p.687823.

[3] Anderson, P.W., 1972. More Is Different: Broken symmetry and the nature of the hierarchical structure of science. Science, 177(4047), pp.393–396.

[4] Ouellette, N.T., 2022. A physics perspective on collective animal behavior. Physical Biology, 19(2), p.021004.

[5] Vicsek, Tamás, and Anna Zafeiris. ”Collective motion.” Physics reports 517, no. 3-4 (2012): 71–140.

[6] Nathan, R., Getz, W.M., Revilla, E., Holyoak, M., Kadmon, R., Saltz, D. and Smouse, P.E., 2008. A movement ecology paradigm for unifying organismal movement research. Proceedings of the National Academy of Sciences, 105(49), pp.19052–19059.

[7] Lewis, M.A., Fagan, W.F., Auger-Methe, M., Frair, J., Fryxell, J.M., Gros, C., Gurarie, E., Healy, S.D. and Merkle, J.A., 2021. Learning and animal movement. Frontiers in Ecology and Evolution, 9, p.681704.

[8] Vicsek, T., Cziŕok, A., Ben-Jacob, E., Cohen, I. and Shochet, O., 1995. Novel type of phase transition in a system of self-driven particles. Physical review letters, 75(6), p.1226.

[9] Romanczuk, P., Couzin, I.D. and Schimansky-Geier, L., 2009. Collective motion due to individual escape and pursuit response. Physical Review Letters, 102(1), p.010602.

[10] Grossman, D., Aranson, I.S. and Jacob, E.B., 2008. Emergence of agent swarm migration and vortex formation through inelastic collisions. New Journal of Physics, 10(2), p.023036.

[11] Peruani, F., Deutsch, A. and Bär, M., 2006. Nonequilibrium clustering of self-propelled rods. Physical Review E, 74(3), p.030904.

[12] Szabo, B., Szöllösi, G.J., Gönci, B., Juŕanyi, Z., Selmeczi, D. and Vicsek, T., 2006. Phase transition in the collective migration of tissue cells: experiment and model. Physical Review E, 74(6), p.061908.

[13] Henkes, S., Fily, Y. and Marchetti, M.C., 2011. Active jamming: Self-propelled soft particles at high density. Physical Review E, 84(4), p.040301.

[14] D’Orsogna, M.R., Chuang, Y.L., Bertozzi, A.L. and Chayes, L.S., 2006. Self-propelled particles with soft-core interactions: patterns, stability, and collapse. Physical review letters, 96(10), p.104302.

[15] Menzel, A.M. and Ohta, T., 2012. Soft deformable self-propelled particles. Europhysics Letters, 99(5), p.58001.

[16] Mikhailov, A.S. and Zanette, D.H., 1999. Noise-induced breakdown of coherent collective motion in swarms. Physical Review E, 60(4), p.4571.

[17] Erdmann, U., Ebeling, W. and Mikhailov, A.S., 2005. Noise-induced transition from translational to rotational motion of swarms. Physical Review E, 71(5), p.051904.

[18] Strömbom, D., 2011. Collective motion from local attraction. Journal of theoretical biology, 283(1), pp.145–151.

[19] Ferrante, E., Turgut, A.E., Dorigo, M. and Huepe, C., 2013. Elasticity-based mechanism for the collective motion of self-propelled particles with springlike interactions: a model system for natural and artificial swarms. Physical review letters, 111(26), p.268302.

[20] Ferrante, E., Turgut, A.E., Dorigo, M. and Huepe, C., 2013. Collective motion dynamics of active solids and active crystals. New Journal of Physics, 15(9), p.095011.

[21] Ginelli, F., Peruani, F., Bär, M. and Chaté, H., 2010. Large-scale collective properties of self-propelled rods. Physical review letters, 104(18), p.184502.

[22] Strandburg-Peshkin, A., Twomey, C.R., Bode, N.W., Kao, A.B., Katz, Y., Ioannou, C.C., Rosenthal, S.B., Torney, C.J., Wu, H.S., Levin, S.A. and Couzin, I.D., 2013. Visual sensory networks and effective information transfer in animal groups. Current Biology, 23(17), pp.R709–R711.

[23] Bleichman, I., Yadav, P. and Ayali, A., 2023. Visual processing and collective motion-related decision-making in desert locusts. Proceedings of the Royal Society B, 290(1991), p.20221862.

[24] Gautrais, J., Ginelli, F., Fournier, R., Blanco, S., Soria, M., Chaté, H. and Theraulaz, G., 2012. Deciphering interactions in moving animal groups.

[25] Strandburg-Peshkin, A., Farine, D.R., Crofoot, M.C. and Couzin, I.D., 2017. Habitat and social factors shape individual decisions and emergent group structure during baboon collective movement. elife, 6, p.e19505.

[26] Lemasson, B.H., Anderson, J.J. and Goodwin, R.A., 2009. Collective motion in animal groups from a neurobiological perspective: the adaptive benefits of dynamic sensory loads and selective attention. Journal of theoretical biology, 261(4), pp.501–510.

[27] Lemasson, B.H., Anderson, J.J. and Goodwin, R.A., 2013. Motion-guided attention promotes adaptive communications during social navigation. Proceedings of the Royal Society B: Biological Sciences, 280(1754), p.20122003.

[28] Ito, S. and Uchida, N., 2024. Selective decision making and collective behavior of fish by the motion of visual attention. arXiv preprint arXiv:2402.09073.

[29] Young, Z. and La, H.M., 2020. Consensus, cooperative learning, and flocking for multiagent predator avoidance. International Journal of Advanced Robotic Systems, 17(5), p.1729881420960342.

[30] Durve, M., Peruani, F. and Celani, A., 2020. Learning to flock through reinforcement. Physical Review E, 102(1), p.012601.

[31] Ĺopez-Incera, A., Ried, K., Müller, T. and Briegel, H.J., 2020. Development of swarm behavior in artificial learning agents that adapt to different foraging environments. PLoS One, 15(12), p.e0243628.

[32] Krongauz, D.L. and Lazebnik, T., 2023. Collective evolution learning model for vision-based collective motion with collision avoidance. PLoS One, 18(5), p.e0270318.

[33] Heins, C., Millidge, B., Da Costa, L., Mann, R.P., Friston, K.J. and Couzin, I.D., 2024. Collective behavior from surprise minimization. Proceedings of the National Academy of Sciences, 121(17), p.e2320239121.

[34] Kim, S.S., Rouault, H., Druckmann, S. and Jayaraman, V., 2017. Ring attractor dynamics in the Drosophila central brain. Science, 356(6340), pp.849–853.

[35] Seelig, J.D. and Jayaraman, V., 2015. Neural dynamics for landmark orientation and angular path integration. Nature, 521(7551), pp.186–191.

[36] Petrucco, L., Lavian, H., Wu, Y.K., Svara, F., S^̌^tih, V. and Portugues, R., 2023. Neural dynamics and architecture of the heading direction circuit in zebrafish. Nature neuroscience, 26(5), pp.765–773.

[37] Sarel, Ayelet, Arseny Finkelstein, Liora Las, and Nachum Ulanovsky. ”Vectorial representation of spatial goals in the hippocampus of bats.” Science 355, no. 6321 (2017): 176–180.

[38] Finkelstein, A., Derdikman, D., Rubin, A., Foerster, J.N., Las, L. and Ulanovsky, N., 2015. Three-dimensional head-direction coding in the bat brain. Nature, 517(7533), pp.159–164.

[39] Taube, J.S., Muller, R.U. and Ranck, J.B., 1990. Head-direction cells recorded from the post- subiculum in freely moving rats. II. Effects of environmental manipulations. Journal of Neuroscience, 10(2), pp.436–447.

[40] Sridhar, V.H., Li, L., Gorbonos, D., Nagy, M., Schell, B.R., Sorochkin, T., Gov, N.S. and Couzin, I.D., 2021. The geometry of decision-making in individuals and collectives. Proceedings of the National Academy of Sciences, 118(50), p.e2102157118.

[41] Poulter, S., Hartley, T. and Lever, C., 2018. The neurobiology of mammalian navigation. Current Biology, 28(17), pp.R1023–R1042.

[42] Kutschireiter, A., Basnak, M.A., Wilson, R.I. and Drugowitsch, J., 2023. Bayesian inference in ring attractor networks. Proceedings of the National Academy of Sciences, 120(9), p.e2210622120.

[43] Stringer, S.M., Trappenberg, T.P., Rolls, E.T. and Araujo, I., 2002. Self-organizing continuous attractor networks and path integration: one-dimensional models of head direction cells. Network: Computation in Neural Systems, 13(2), pp.217–242.

[44] York, L.C. and van Rossum, M.C., 2009. Recurrent networks with short term synaptic depression. Journal of computational neuroscience, 27(3), pp.607–620.

[45] Jeffery, K.J., Page, H.J. and Stringer, S.M., 2016. Optimal cue combination and landmark- stability learning in the head direction system. The Journal of physiology, 594(22), pp.6527–6534.

[46] Stringer, S.M., Rolls, E.T. and Trappenberg, T.P., 2004. Self-organising continuous attractor networks with multiple activity packets, and the representation of space. Neural networks, 17(1), pp.5–27.

[47] Ocko, S.A., Hardcastle, K., Giocomo, L.M. and Ganguli, S., 2018. Emergent elasticity in the neural code for space. Proceedings of the National Academy of Sciences, 115(50), pp.E11798–E11806.

[48] Vafidis, P., Owald, D., D’Albis, T. and Kempter, R., 2022. Learning accurate path integration in ring attractor models of the head direction system. Elife, 11, p.e69841.

[49] Robinson, B.S., Norman-Tenazas, R., Cervantes, M., Symonette, D., Johnson, E.C., Joyce, J., Rivlin, P.K., Hwang, G.M., Zhang, K. and Gray-Roncal, W., 2022. Online learning for orientation estimation during translation in an insect ring attractor network. Scientific reports, 12(1), p.3210.

[50] Stringer, S.M., Rolls, E.T. and Trappenberg, T.P., 2005. Self-organizing continuous attractor network models of hippocampal spatial view cells. Neurobiology of learning and memory, 83(1), pp.79–92.

[51] Wang, R. and Kang, L., 2022. Multiple bumps can enhance robustness to noise in continuous attractor networks. PLOS Computational Biology, 18(10), p.e1010547.

[52] Gorbonos, D., Gov, N.S. and Couzin, I.D., 2024. Geometrical Structure of Bifurcations during Spatial Decision-Making. PRX Life, 2(1), p.013008.

[53] Oscar, L., Li, L., Gorbonos, D., Couzin, I.D. and Gov, N.S., 2023. A simple cognitive model explains movement decisions in zebrafish while following leaders. Physical Biology, 20(4), p.045002.

[54] Katz, Y., Tunstrøm, K., Ioannou, C.C., Huepe, C. and Couzin, I.D., 2011. Inferring the structure and dynamics of interactions in schooling fish. Proceedings of the National Academy of Sciences, 108(46), pp.18720–18725.

[55] Herbert-Read, J.E., Perna, A., Mann, R.P., Schaerf, T.M., Sumpter, D.J. and Ward, A.J., 2011. Inferring the rules of interaction of shoaling fish. Proceedings of the National Academy of Sciences, 108(46), pp.18726–18731.

[56] Couzin, I.D., Krause, J., Franks, N.R. and Levin, S.A., 2005. Effective leadership and decision-making in animal groups on the move. Nature, 433(7025), pp.513–516.

[57] Mussells Pires, P., Zhang, L., Parache, V., Abbott, L.F. and Maimon, G., 2024. Converting an allocentric goal into an egocentric steering signal. Nature, pp.1–11.

[58] Suthana, N.A., Ekstrom, A.D., Moshirvaziri, S., Knowlton, B. and Bookheimer, S.Y., 2009. Human hippocampal CA1 involvement during allocentric encoding of spatial information. Journal of Neuroscience, 29(34), pp.10512–10519.

[59] Klatzky, R.L., 1998. Allocentric and egocentric spatial representations: Definitions, distinctions, and interconnections. In Spatial cognition: An interdisciplinary approach to representing and processing spatial knowledge (pp. 1–17). Berlin, Heidelberg: Springer Berlin Heidelberg.

[60] Li, D., Karnath, H.O. and Rorden, C., 2014. Egocentric representations of space co-exist with allocentric representations: evidence from spatial neglect. Cortex, 58, pp.161–169.

[61] McNaughton, B.L., Battaglia, F.P., Jensen, O., Moser, E.I. and Moser, M.B., 2006. Path integration and the neural basis of the’cognitive map’. Nature Reviews Neuroscience, 7(8), pp.663–678.

[62] Filimon, F., 2015. Are all spatial reference frames egocentric? Reinterpreting evidence for allocentric, object-centered, or world-centered reference frames. Frontiers in human neuroscience, 9, p.648.

[63] Kraft, P., Evangelista, C., Dacke, M., Labhart, T. and Srinivasan, M.V., 2011. Honeybee navigation: following routes using polarized-light cues. Philosophical transactions of the royal society B: biological sciences, 366(1565), pp.703–708.

[64] Horváth, G. and Varjú, D., 2004. Polarized light in animal vision: polarization patterns in nature. Springer Science & Business Media.

[65] Wiltschko, R. and Wiltschko, W., 2015. Avian navigation: a combination of innate and learned mechanisms. Adv. Study Behav, 47, pp.229–310.

[66] Wiltschko, R. and Wiltschko, W., 2023. Animal navigation: how animals use environmental factors to find their way. The European Physical Journal Special Topics, 232(2), pp.237–252.

[67] Mouritsen, H., 2018. Long-distance navigation and magnetoreception in migratory animals. Nature, 558(7708), pp.50–59.

[68] Hopfield, J.J., 1982. Neural networks and physical systems with emergent collective computational abilities. Proceedings of the national academy of sciences, 79(8), pp.2554–2558.

[69] Green, R.F., 1984. Stopping rules for optimal foragers. The American Naturalist, 123(1), pp.30–43.

[70] McCall, J.J. and Lippman, S.A., 1984. Ecological decision making and optimal stopping rules (No. 189). Diskussionsbeiträge-Serie A.

[71] Romey, W.L., Smith, A.L. and Buhl, C., 2015. Flash expansion and the repulsive herd. Animal Behaviour, 110, pp.171–178.

[72] Romey, W.L. and Lamb, A.R., 2015. Flash expansion threshold in whirligig swarms. PLoS One, 10(8), p.e0136467.

[73] Couzin, I.D. and Krause, J., 2003. Self-organization and collective behavior in vertebrates. Advances in the Study of Behavior, 32(1), pp.10–1016.

[74] Salahshour, M., 2019. Phase diagram and optimal information use in a collective sensing system. Physical review letters, 123(6), p.068101.

[75] Joshi, P.S., 1994. Global aspects in gravitation and cosmology. Oxford University Press.

[76] Amit, D.J. and Amit, D.J., 1989. Modeling brain function: The world of attractor neural net- works. Cambridge university press.

